# Unexpected Early Proteomic Changes in Alzheimer’s Disease Model Mice Synaptosomes

**DOI:** 10.1101/144972

**Authors:** Kerri Ball, Addolorata Pisconti, Kelly Grounds, William M. Old, Michael H. B. Stowell

## Abstract

We have employed label-free quantitative proteomics of wild-type and Alzheimer’s disease (AD) model mice synaptosomes to investigate proteomic changes occurring during AD progression as a prelude to analysis in humans. More than 4000 proteins were analyzed using multiple analysis tools and statistical criteria. Pathway enrichment identified numerous pathways consistent with the current AD knowledge base, including dysregulation of Glutamate Receptor Signaling, Synaptic Long Term Potentiation and Depression, Rho and Rac Signaling, Calcium Signaling, and Oxidative Phosphorylation and Mitochondrial Dysfunction. Additionally, the data demonstrate that a large number of changes occur in the proteome very early relative to the onset of both traditional disease markers such as amyloid accumulation, tau phosphorylation and cognitive dysfunction. These early changes include a number of dysregulated proteins that have novel associations with AD progression. These results reinforce the importance of mechanistic investigations in early disease progression long before the classical markers of Alzheimer’s disease are observed.

## Introduction

The synapse is the localized contact between nerve cells required for signal transmission and AD is considered by many to be a synaptic disease. This cell-to-cell communication is characterized by complex protein-driven molecular mechanisms including synthesis, delivery, storage, docking, fusion, neurotransmitter release and reuptake (1). Synapses can be studied by isolation of synaptosomes which contain the complete presynaptic terminal, including mitochondria and synaptic vesicles, along with the postsynaptic membrane and the postsynaptic density. Several proteomic studies of synaptosomes have previously been performed (2–5). However, only recently has mass spectrometric analysis reached the level of technical advancement necessary for a direct and comprehensive analysis of the synaptic proteome (6). These advances in proteomics technologies allow direct and unbiased examination of protein level differences in neurodegenerative diseases and have great potential to shed new light on disease pathogenesis. Here, we employed these technical advancements in mass spectrometry for the detection of more than 4,000 synaptosomal proteins using label-free quantitative proteomics to characterize the proteome changes that occur in Alzheimer’s disease (AD) model mice. Multiple structural and/or metabolic proteins have been reported to have altered expression in AD supporting a high depth quantitative proteomic analysis for target discovery (7–10).

## Results

### Mouse Model Characterization

The B6C3-Tg(APPswe,PSEN1dE9)85Dbo/Mmjax mouse model (Tg-AD) of Alzheimer’s disease is a widely used model for AD; it contains a chimeric mouse/human amyloid precursor protein (APP)(Mo/HuAPP695swe) and human presenilin 1 (PS1-dE9); both driven by the prion protein promoter and therefore expressed in central nervous system neurons (11–15). These two insertions favor processing through the β-secretase pathway and, thus, elevate the amount of amyloid-beta (Aβ) fragments produced from the APP transgene. To validate accumulation of Aβ fragments in transgenic mice, we used commercially available ELISA kits specific for Aβ 1-42. As shown in Fig1, both Aβ 1-42 accumulates in the brain of Tg-AD mice in an age dependent manner, supporting the choice of this mouse strain as a relevant model of AD.

**Fig 1.**
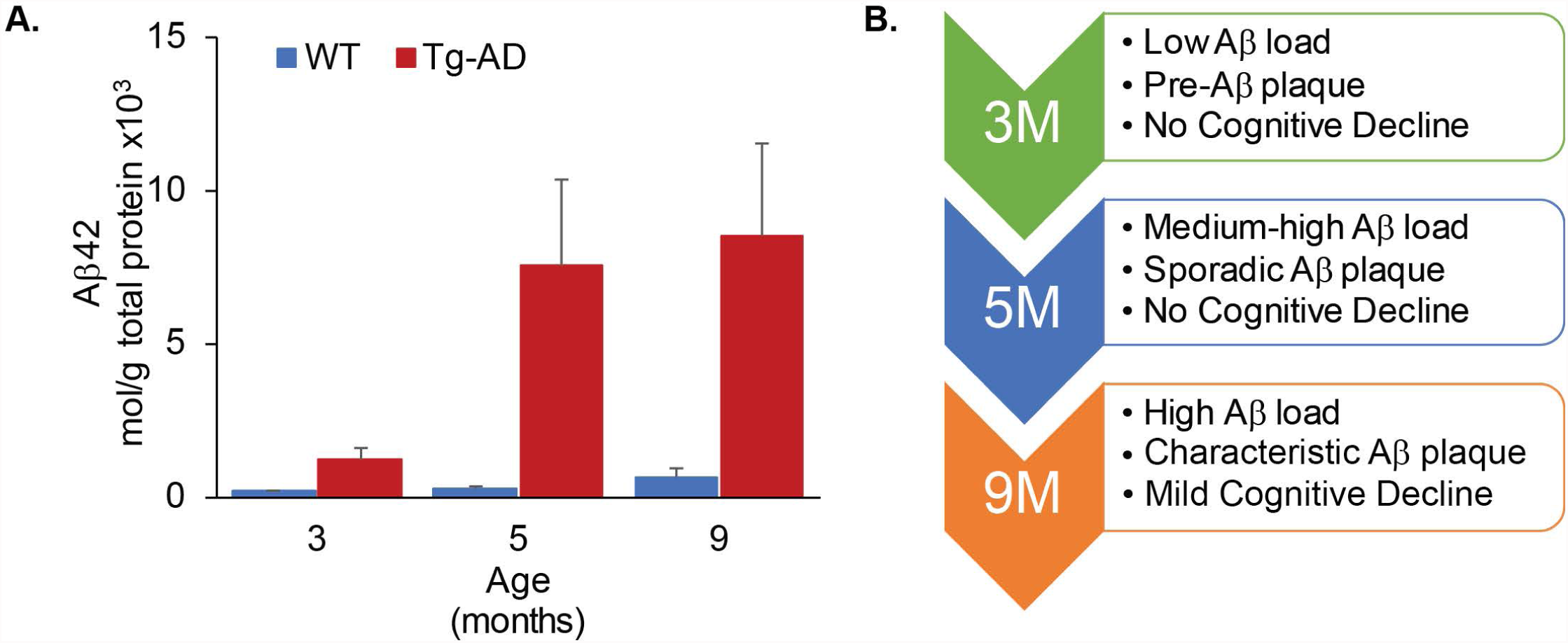
Amyloid accumulation in the APPswe/PSEN1dE9 Alzheimer’s disease mouse model. (A.) Human Aβ42 was quantitated using commercially available sandwich ELISA kits at the indicated ages using both wild-type (WT) and Tg-AD (B6C3-Tg (APPswe, PSEN1dE9)85Dbo/J)) mice. Amounts of Aβ42 were normalized to total protein as determined by BCA Assay. (B.) Three, five and nine month old Tg-AD mice were chosen for high depth proteomic analysis based on the stages of Aβ load, plaque development, and cognitive decline.

Based on our Aβ accumulation data (Fig1) and upon previous characterizations of the APPswe, PSEN1dE9 biogenic mouse model which describe the timing of cognitive impairment and plaque formation (11–15), we chose to carry out a proteomic analysis of the synaptosomes of three, five, and nine month-old mice. These three age groups represent three distinct stages of AD, summarized in Fig1C. The three month-old Tg-AD age group have minimal accumulation of Aβ1-42, normal cognitive function, and complete absence of plaques. The five month-old Tg-AD mice have relatively high levels of Aβ1-42 which are accompanied by the presence of sporadic plaques although no cognitive decline has been reported. In contrast, the nine month-old Tg-AD mice represent the post-plaque and post-cognitive decline stage of AD.

### Workflow

Synaptosomes from 3, 5, and 9 month-old Tg-AD and wild-type (WT) mice were isolated according to standard protocols (16). Filter Assisted Protein Preparation (FASP) was utilized to obtain pure peptides for LC-MS/MS analysis (17). Peptide samples were analyzed by online two dimensional reverse phase (RP/RP) nanoflow HPLC-MS/MS. Raw mass spectrometry data was converted to protein abundance using chromatography feature finding software MaxQuant (version 1.5.2.8) (18–20).

A total of 4,655 proteins were identified across all 18 samples using MaxQuant (18–20). All proteins categorized as potential contaminants, reverse sequence, and/or only identified by site were removed from the analysis. Biological replicates were than categorically annotated into 6 groups and proteins containing less than 2 valid values in each group were removed from the analysis, thus reducing the matrix to 3312 quantifiable proteins. (S1 Data File).

**S1 Data File. Protein Quantification and Analysis.**

### Qualitative Analysis

A qualitative evaluation of the MS data was performed on both the WT and AD data. While biological variation is expected and accepted among replicates, due to complex and step-wise collection of RP/RP HPLC MS/MS data technical variations should be evaluated. For this analysis, we started with a hierarchical cluster analysis (HCA) performed using complete-linkage clustering with Euclidean distance metrics of the nine WT samples (Fig2A) and the nine AD samples (Fig2B). As shown in Fig2A, the sample WT_3M_12f-C was identified as a potential outlier to other eight WT samples that are all merged into one cluster. The sub-clusters of the eight WT samples include pair-wise clustering of 3M with 5M and 5M with 9M, suggesting that age-dependent difference between the three WT age groups are negligible. In contrast, the main 2 clusters in the AD HCA (Fig2B) have a relatively short distance between them and the sub-clustering pairs include age matched AD samples, such as the pairing of AD_3M_14c-F with AD_3M_12f-AD, AD_9M_16c-CF with AD_9M_16a-D, and AD_5M_11a-C with AD_5M_11a-AB. Thus, the AD HCA suggests that differences between the age groups and, moreover, differences between the three selected stages of Alzheimer’s disease will be observed.

**Fig 2.**
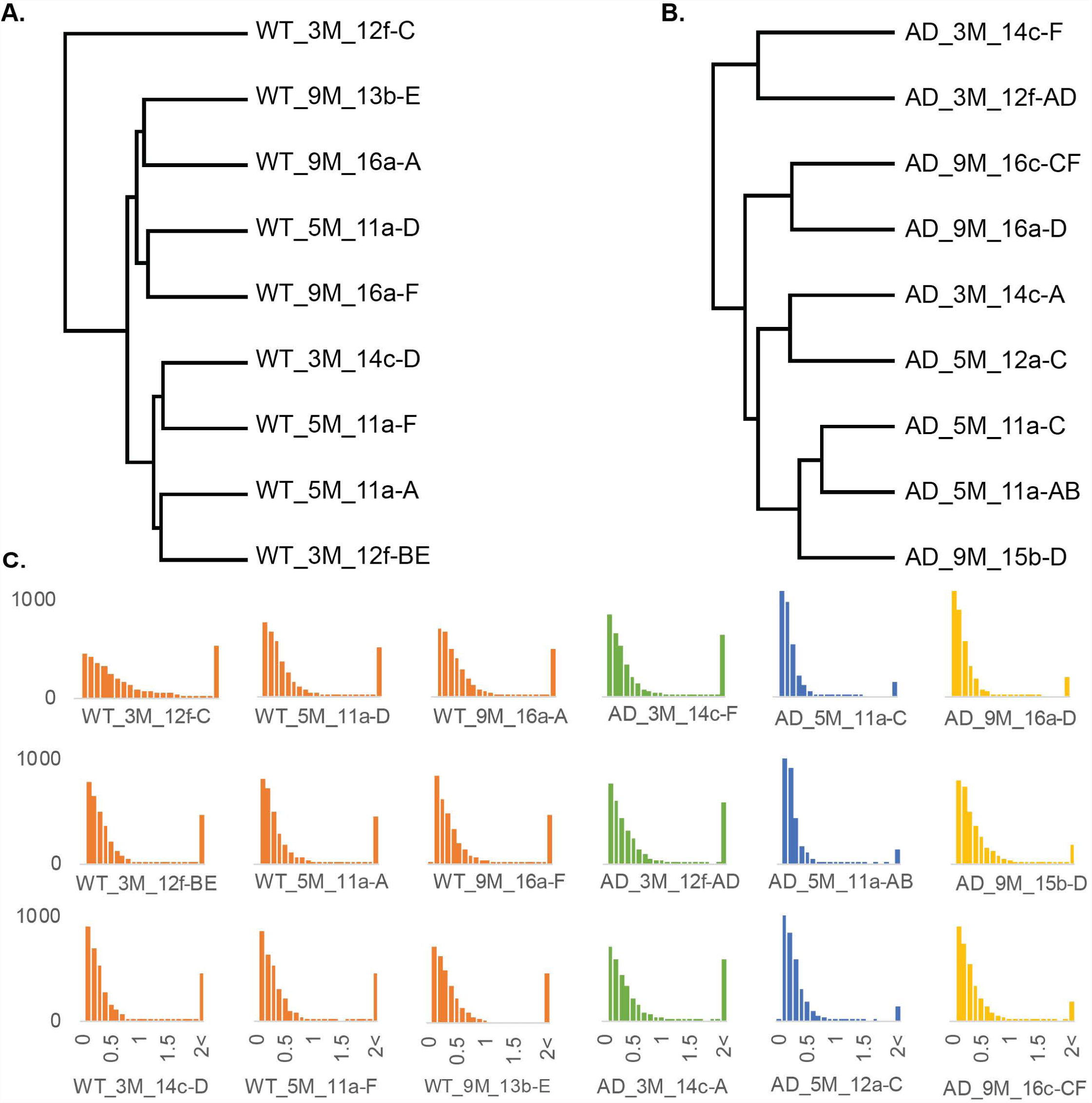
Qualitative Assessment of MS/MS data. A dendogram of the WT samples (A.) and the AD samples (B.) produced using Euclidean distance with complete linkage of LFQ intensity data with all invalid values removed. (B.) Histogram of absolute log_2_ expression values (ORANGE: WT_x – WT_average =, GREEN: 3M_x – 3M_average, BLUE: 5M_x – 5M_average, YELLOW: 9M_x – 9M_average).

To further explore the variability between replicates, we evaluated the log_2_ fold change of individual samples from the average (Fig2C, ORANGE: WT from the average WT, GREEN: 3M_AD from the average 3m Tg-AD, BLUE: 5M_AD from the average 5m Tg-AD, and YELLOW: 9M_AD from the average 9m Tg-AD). The expectation is that all samples included in the average should have a similar shaped distribution, thus the shorter and broader shape of the sample WT_3M_12f-C distribution compared to the other WT samples further supports the identification of this sample as an outlier. The 3M_AD samples all have a similar distribution, despite the Euclidean distance of sample AD_3M_14c-A from the other two 3M_AD samples (Fig2B). While the AD_5M and AD_9M sample distributions are a clear reflection the short Euclidean distances observed in Fig2B; in particular, the distance between AD_5M_11a-C and AD_5M_11a-AB is shorter than for any other pairing and the distribution of these samples is taller and more narrow than any of the other samples.

Together, this qualitative evaluation of the MS/MS data identifies sample WT_3M_12f-C as an outlier that will be excluded from the quantitative analysis. Additionally, this analysis supports that age dependent biological variability between the control groups is very small. Grouping the 8 WT samples in one control group increases the statistical power of the analysis, but may sacrifice age specific variations, thus further evaluation of age dependent protein abundance was analyzed.

To determine if pooling the WT samples would be appropriate, we analyzed potential age-dependent changes in the WT samples using an ANOVA multi-sample test. No statistically significant proteins were identified using a loose False Discover Filter (FDR, Bengamini-Hochburg) α = 0.25. Thus, to identify genes that would likely lose statistical relevance if an average WT was used rather than age matched controls, we applied a p-Value cut-off of 0.1 with no FDR and an absolute log_2_ fold change filter of 0.5. We identified 73 proteins of interest (S1 Data File) that could be lost if we pooled the WT samples into a single control group. However, despite these potential loses, we made the decision to group all of the WT samples into a single control group.

### Quantitative Assessment of Alzheimer’s Disease Progression Proteome

R-limma (21–23) was used to perform a quantitative assessment of proteome differences between the following groups: 3m Tg-AD & WT, 5m Tg-AD & WT, and 9m Tg-AD & WT. Empirical Bayes statistics was used to calculate p-Values. Using a Bengamini-Hochburg FDR α = 0.1, (Fig3; S1 Data File). Surprisingly, the majority of statistically significant proteins identified were only significant in the pre-plaque, 3m Tg-AD, stage of AD when Aβ levels are still very low. The protein expression profiles of the top most dysregulated proteins (log_2_ FC >= |1|) are shown in Fig4. We noted that even though the selection of significant proteins was dominated by the 3m Tg-AD statistical analysis, clear age dependent data trends were observed; we clustered the proteins based on these expression trends. This clustering shows groups of proteins that decrease (clusters 2 & 3) and increase (clusters 8, 9, & 10) during AD progression, that are consistently up (cluster 1 & 2) or down (cluster 10 & 11), and that have stage specific protein dysregulation (clusters 4, 5, 6, & 7). Eleven of the proteins of interest identified have a previous association with Aβ and/or AD (Fig4 *GENE), while 73 of these high confidence proteins (log_2_ FC >= |1|) are novel to our understanding of AD progression.

**Fig 3.**
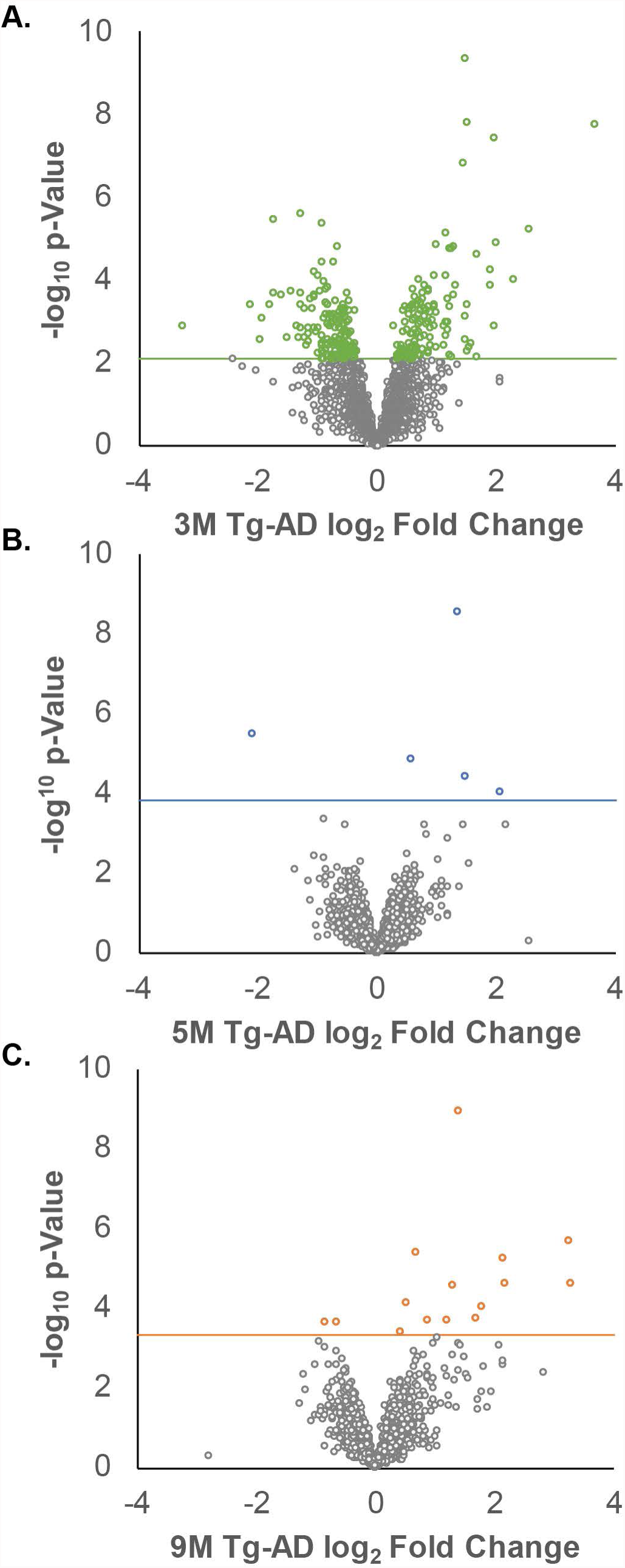
Quantitative Analysis of Age-Dependent Alzheimer’s Disease. R-limma was used to calculate the p-Value and log_2_ fold change for the 3312 quantifiable proteins using the following groupings: (A.) 3m Tg-AD & WT, (B.) 5m Tg-AD & WT, and (C.) 9m Tg-AD & WT. A Bengamini-Hochburg FDR α = 0.1 is indicated with a solid line in each graph and the significant proteins are colored. This data and the corresponding gene names are included in Supplement File 1.

**Fig 4.**
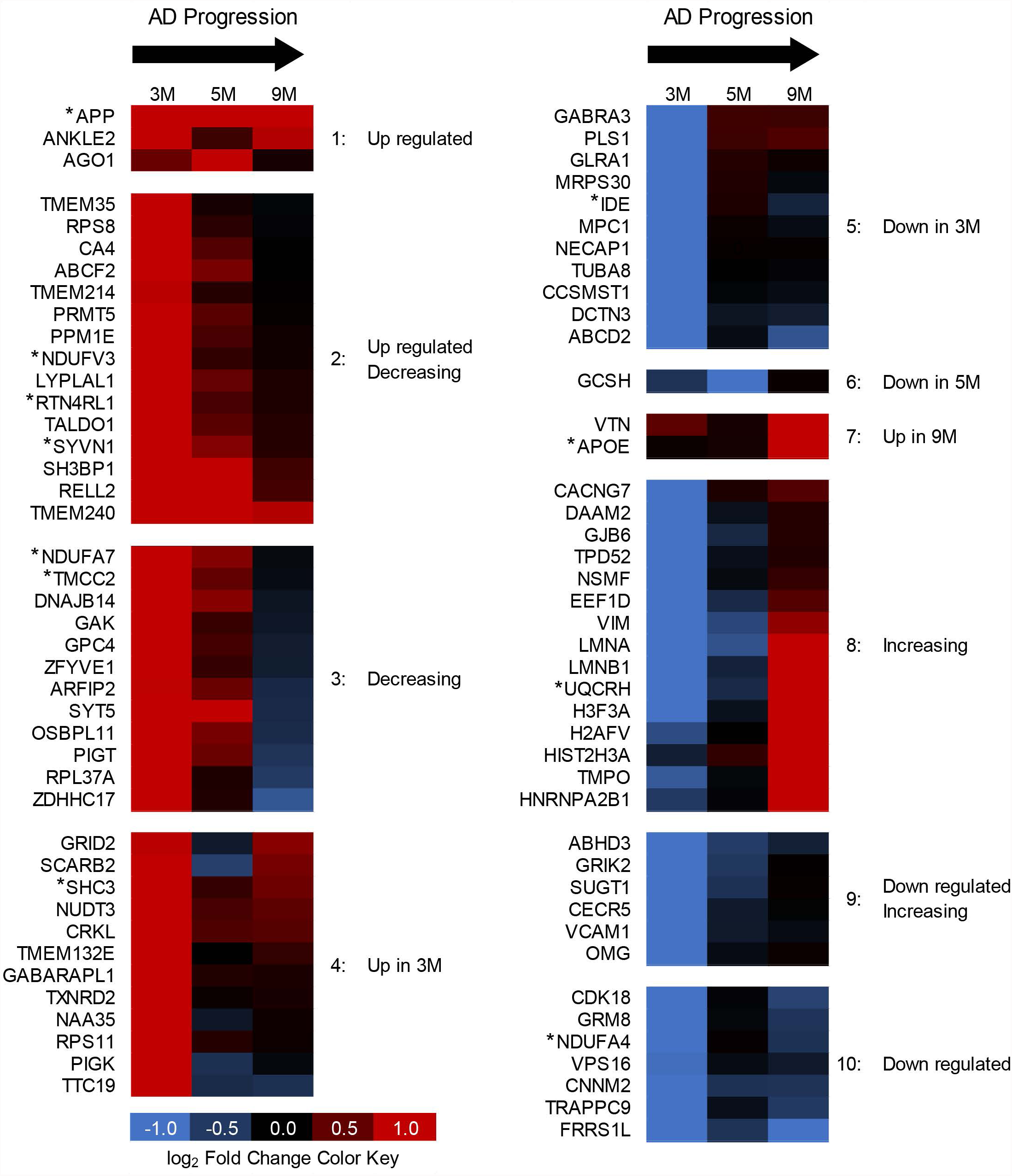
High-Confidence Protein Expression Profiles and Clusters. Proteins were filtered using an FDR = 0.1 and an absolute FC = 1 in at least one age group. Proteins were clustered based on observable FC trend. Proteins with a previously recognized association with Aβ (24) and/or recognized by Kegg as “Alzheimer’s Disease: are indicated by * next to the gene name.

### STEM Analysis

As mentioned above, we observed many proteins that appeared to have a linear trend of protein expression. Time-dependent data trends can further increase confidence in the data, and analysis of these trends may even identify addition statistically significant protein changes. To further explore proteins with trending protein profiles we utilized the Short Time-series Expression Miner (STEM)(25). We used a maximum of 50 model profiles and maximum unit change of 3 (log_2_ FC) between time points to profile the matrix of 3312 quantifiable proteins at the three Tg-AD age groups. STEM profile enrichment identified 121 proteins that follow one of five significant profiles (S2 Data File). Fig5A-E shows these significant STEM profiles (black); the protein expression patterns that were fitted to these profiles are plotted along with the STEM profiles. Not quite half the proteins identified by STEM (58/121 proteins) were identified as significant using the empirical Bayes statistical enrichment, while 63 new proteins were added to the list of proteins dysregulated during the progression of AD.

**Fig 5.**
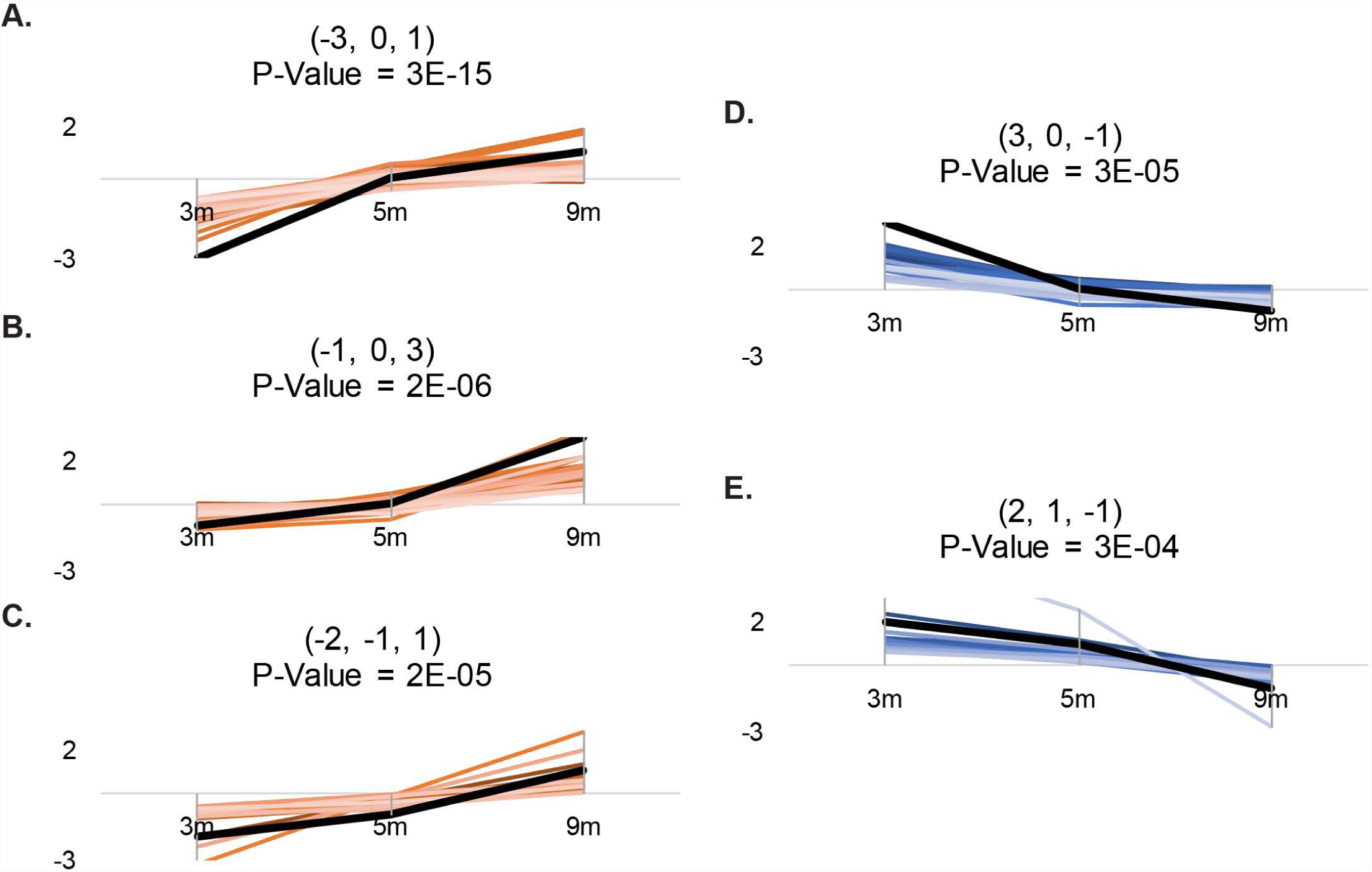
Significant Protein Trends identified by STEM. Five protein expression profiles were identified as having more proteins fitting to the profile than would be expected by random chance. These significant profiles included three increasing profiles with 41 proteins fitting the -3, 0, 1 log_2_ FC profile (A.) in the 3m, 5m, and 9m age groups, respectively, 25 proteins fitting the -1, 0, 3 log_2_ FC profile (B.), and 16 proteins fitting the -2, -1, 1 log_2_ FC profile (C.). Two decreasing profiles were also identified; these were characterized by 25 proteins fitting the 3, 0, -1 log_2_ FC profile (D.) and 14 proteins fitting the 2, 1, -1 profile (E.) The profile summaries exported from STEM can be found in S2 Data File and the protein-to-profile assignments can be found in S2 Data File as well as in S1 Data File.

**S2 Data File. Time-series expression analysis by STEM.**

### Canonical Pathway Enrichment

The benefit of adding proteins identified by STEM is demonstrated in Fig6 where Canonical Pathway Enrichment (by Ingenuity Pathway Analysis, IPA) is compared between three data filters for the 3m Tg-AD data set: a strict filter (FDR 0.1 + log_2_ FC >= |1|), an FDR only filter (FDR 0.1), and a STEM enriched filtered data set (FDR 0.1 + log_2_ FC >= |0.4|, +STEM proteins) for a select group of canonical pathways (complete list: S3 Data File). P-values for canonical pathways are calculated in IPA using a right-tailed Fisher Exact Test that considers the overlap of observed and predicted genes in a pathway. Thus, while strict data filters will result in a high confidence gene list (Fig4), a short gene list will also produce a low confidence canonical pathway enrichment. Additionally, Fig6 & S3 Data File show that, with few exceptions, expanding the gene list to include lower confidence protein expressions identified using STEM adds to our pathway confidence (p-Value) or in other words, “what is significant becomes MORE significant”. IPA also assigns a Z-score that assesses the match of observed and predicted up/down regulation patterns. Therefore, the age-dependent FC for all proteins in the Canonical Pathway are considered. In IPA, a Z-score greater than 2 or less than -2 is considered predictive; positive Z-scores indicate activation and negative Z-scores indicate repression of the described function. As shown in Fig6, the STEM enriched protein list allows higher confidence directional predictions in comparison the FDR only filtered protein list. No predictable directionality was found in any of the enriched Canonical Pathways when the high confidence protein list (FDR 0.1, |FC|>1) was used.

**Fig 6.**
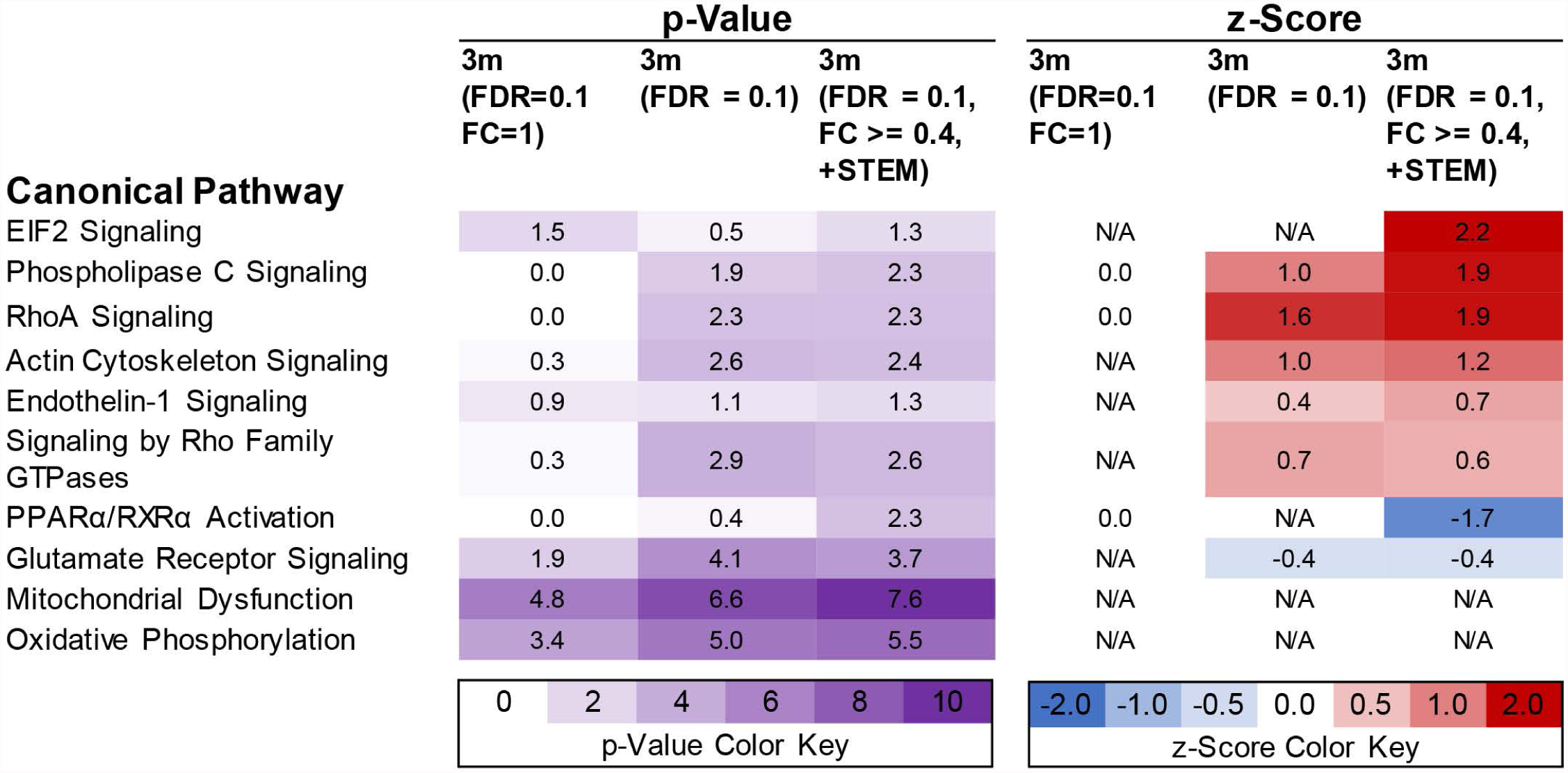
STEM proteins add confidence and direction to Canonical Pathway Enrichment analysis. Ingenuity Pathway Analysis (IPA) was used to analyze the 3m Tg-AD Canonical Pathway Enrichment using three different data filters: the high confidence protein list from Fig4, a standard filter (FDR = 0.1 only), and the STEM expanded FDR = 0.1 and absolute FC >= 0.4. p-Values and z-scores as calculated by IPA are shown for the top Canonical Pathways. The complete Canonical Pathway Enrichment list can be found in S3 Data File.

**S3 Data File. Canonical Pathway Enrichment.**

With this in mind we utilized the protein expression data from all proteins identified as significant (FDR = 0.1, absolute log_2_ FC >= 0.4) plus the STEM identified proteins to run a canonical pathway enrichment analysis. All pathways with a z-score in at least one age group are shown in Fig7. Z-scores greater than or equal to the absolute value of two are considered predictive. All dysregulated pathways, including those without a z-score, can be found in S3 Data File.

**Fig 7.**
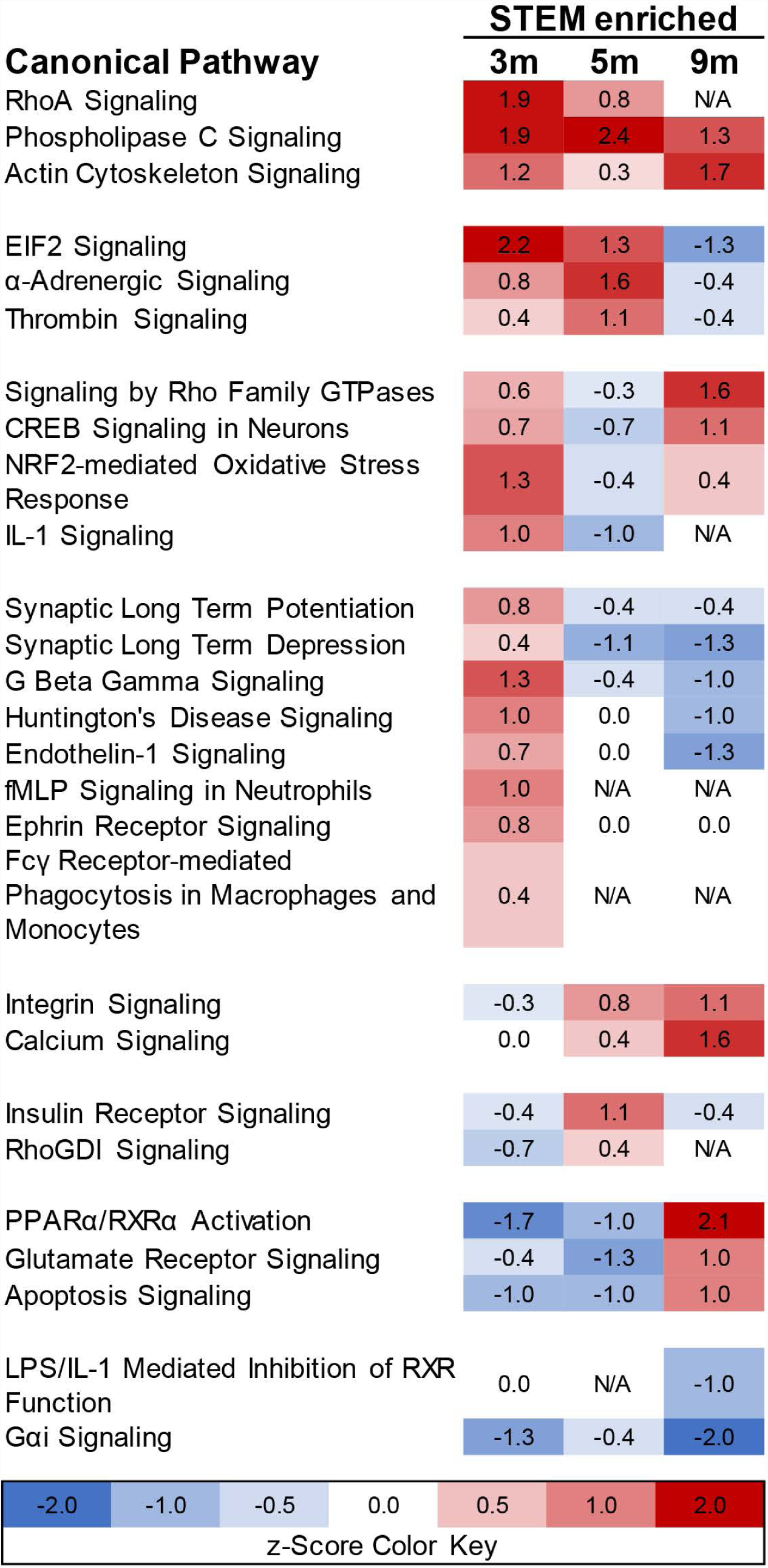
Ingenuity Pathway Analysis (IPA) predicted direction of Canonical Pathways. IPA was used to analyze compare the directionality (z-scores) of Canonical Pathways identified enriched in the STEM expanded FDR = 0.1 and absolute FC >= 0.4 protein list. Canonical pathways with valid/non-zero z-score in at least one age group are shown here. The complete Canonical Pathway Enrichment list can be found in S3 Data File.

## Discussion

The most well-known and well-studied histopathological hallmark of AD is an increase in Aβ peptide abundance and subsequent formation of amyloid plaques. Accordingly, the mouse model used in our studies should provide insights in downstream mechanisms following Aβ production. Aβ is believed to be a crucial pathogenic factor in AD development. Recent evidence indicates that the soluble-oligomeric forms of Aβ are primarily responsible for the neurodegeneration and loss of synaptic function characteristic of later stages of AD, and this soluble-Aβ hypothesis is further supported by recent clinical data on aducanumab, a human monoclonal antibody shown to reduce soluble and insoluble Aβ (26).

Surprisingly, we observed that the largest changes to the synaptosomal proteome occurred in the 3m Tg-AD mouse model where Aβ levels are relatively low, Aβ plaques are absent, and no cognitive decline is observable. While this is consistent with soluble-oligomeric forms of Aβ being primarily responsible for the pathogenesis of AD, it is notable that despite the large proteomic changes, the 3m Tg-AD mice have no observable phenotype. Very recently, a new AD phenotype has been observed: olfactory recall impairment occurs up to ten years prior to the onset of cognitive decline (27,28). This suggests that the observed proteomic changes in the 3m Tg-AD mouse may have correlated phenotypes that remain to be identified. However, it could also suggest that Aβ has a significant biological impact on the brain, even at low levels, and that compensatory mechanisms are not only active but, most importantly, are biological efficacious during the preclinical stages of AD. Although the ultimate progression of AD suggests that any early neuroprotective response to Aβ is not sustainable or is not sufficient to prevent neurotoxicity.

In typical AD progression, Aβ promotes disturbances in a number of pathways that ultimately lead to neurotoxicity. Specifically, Aβ oligomers have been reported to induce NMDA receptor activation, mitochondrial Ca2+ overload/membrane depolarization, oxidative stress and apoptotic cell death (29–32). Consistent with this pathology, we observed dysregulation in these Canonical Pathways: Glutamate Receptor Signaling, Synaptic Long Term Potentiation and Depression, Calcium Signaling, and Oxidative Phosphorylation and Mitochondrial Dysfunction (Figures 6 & 7, S3 Data File). Beyond observing dysregulation of Canonical Pathways that are consistent with what is already known about AD, we identified 73 proteins with high confidence (log_2_ FC >= |1|, Fig4) that are novel in our understanding of AD progression, but that are consistent with previous studies. While refraining from going through all proteins/pathways, a striking example of how our data supports the existing knowledge can be found by close inspection of the RhoA Pathway depicted in Fig8. Specifically, our data shows an increase RhoA GEF protein (ARHGEF1) and a decrease in RhoA GAP protein (RHOGAP) in the 3m Tg-AD (Fig8A), consistent with activation of RhoA Signaling Pathway (Fig8B). RhoA activation, as shown in Fig8A & B, leads to the activation a number of kinases whose downstream activities regulate the actin cytoskeleton. ROCK, for example, is a kinase downstream of RhoA that activates LIMK which in turn phosphorylates, and thus inhibits, the actin severing protein, cofilin (Fig8). However, in direct opposition to the IPA’s predicted state of cofilin, previous observations with Aβ1-42 treatment show an increase in dephosphorylated cofilin (33) and an increase of cofilin translocation into the mitochondria (consistent with dephosphorylated cofilin) (34). Moreover, in AD patients, a loss or shortening of dendritic spines is observed, consistent with loss of cytoskeletal stability. Previous studies have also shown that Aβ1-42 treatment of hippocampal neurons induced increased activity in Rac1, Cdc42, and PAK1 (33), which, like RhoA, are also involved in activation of LIMK and the phosphorylation of cofilin. Together, this data suggests that Aβ either inhibits phosphorylation or promotes dephosphorylation of cofilin and that activation of upstream activators of LIMK may be a compensation mechanism for the increase in dephosphorylated cofilin. Potential mechanisms of Aβ’s regulatory role on cofilin phosphorylation and dephosphorylation are illustrated in Fig9A.

**Fig 8.**
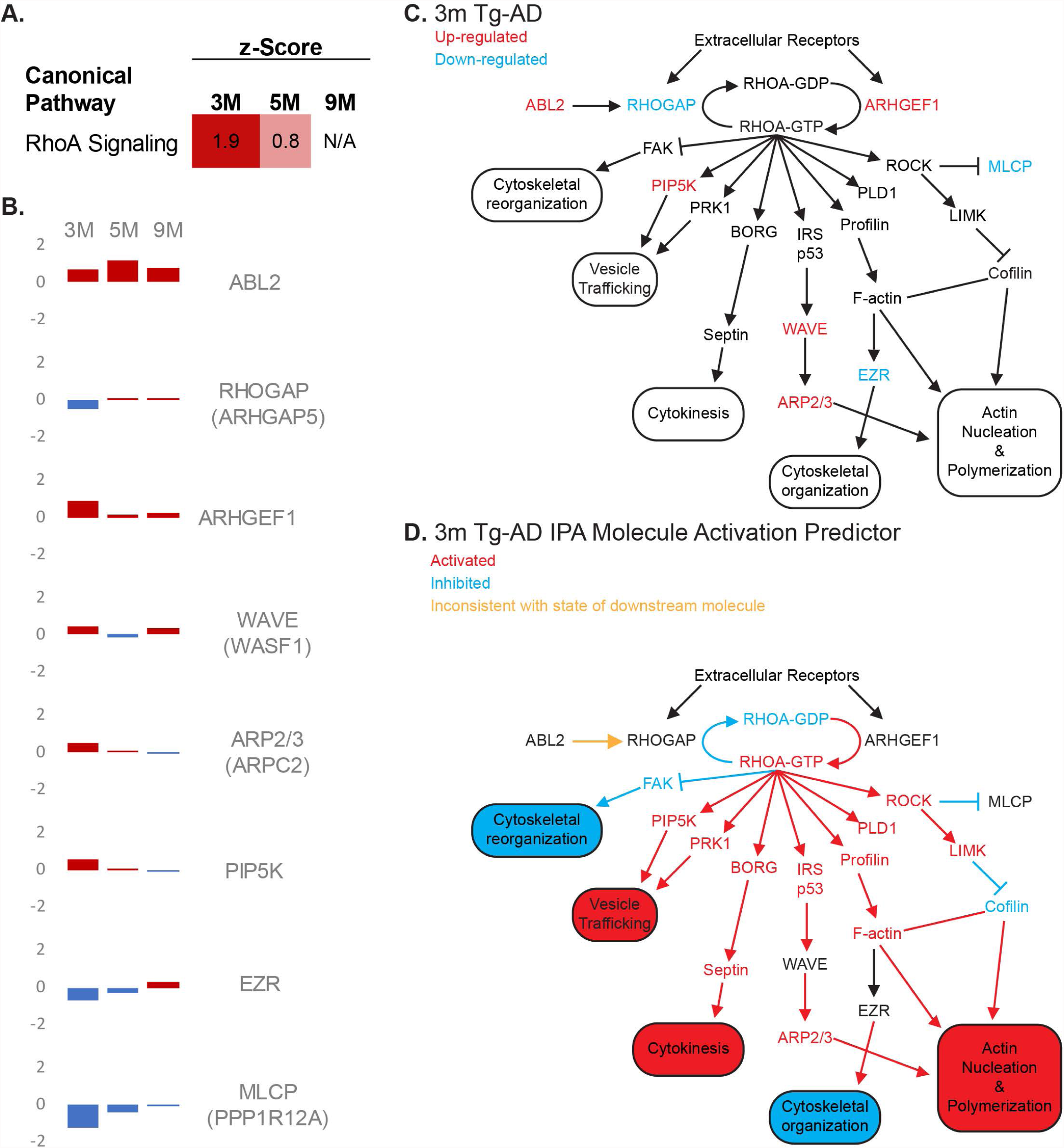
RhoA Signaling Pathway activation in early AD. (A.) Canonical Pathway enrichment analysis identified RhoA as activated in the 3m Tg-AD. (B.) The log_2_ FC of eight proteins within the RhoA Signaling Pathway were used to calculate the z-score and the predicted directionality of RhoA Signaling. (C.) These molecules are colored red (up-regulated) or blue (down-regulated) in a schematic of the RhoA Signaling pathway in the 3m Tg-AD. (D.) IPA’s Molecular Activation Predictor illustrates the predicted outcome of the 3m Tg-AD protein abundance changes. (Red = activated; Blue = inhibited; Yellow = inconsistent with state of downstream molecule)

**Fig 9.**
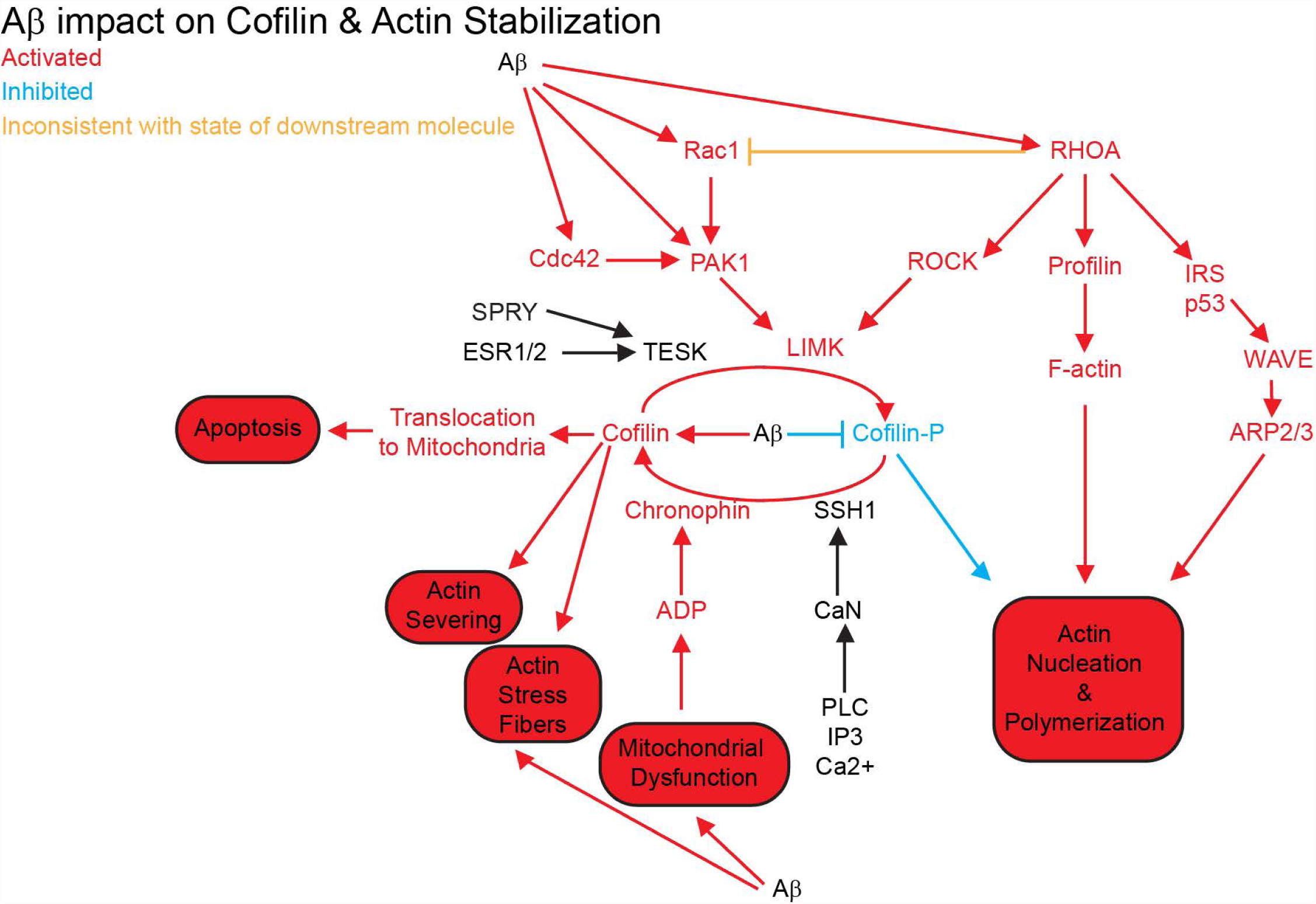
Proposed Aβ’s impact on Cofilin Phosphorylation and actin stabilization. Our study, in concert with previous studies, implicates Aβ involvement in the activation of a number of molecules upstream of Cofilin regulation and actin stabilization. (Red = activated; Blue = inhibited; Yellow = inconsistent with state of downstream molecule)

Compensation for Aβ induced dysregulation of cofilin-actin dynamics would be required for sustained neural function and survival. Cofilin-actin regulation is critical for morphogenesis and the structural dynamics of neural spines and has been strongly implicated in synaptic trafficking of AMPA receptors during Synaptic Potentiation and Depression (35,36). In the 3m Tg-AD, we do not see a significant difference in the number of AMPA receptors (GRIA). However, we assume there is successful compensatory stabilization of actin filaments through the RhoA Signaling Pathway.

The RhoA Signaling Pathway represents only one of many pathways that were found to be dysregulated in the pre-clinical 3m Tg-AD model. Understanding how these pathways take part in the response to Aβ may lead to new therapeutic avenues for AD. Additionally, recognition of the early dysregulation of these pathways may help identify new, pre-clinical, phenotypes for AD.

## Methods

### Mice

Mice used in the experiments were housed in accordance with protocols approved by the Institutional Animal Care and Use Committee at University of Colorado and all experiments were conducted according to the NIH Guide for the Care and Use of Laboratory Animals. The B6C3-Tg(APPswe, PSEN1dE9)85Dbo/J) breeding pair was obtained from The Jackson Laboratory (stock number 004462). Colony was maintained using +/+ sibling x Hemi zygote. Ear punches were taken at approximately 10 days old and PCR identification was performed to identify AD transgenic mice and the non-transgenic littermate controls. The AD transgenic mice and the non-transgenic littermate controls of the same sex were housed in individual ventilated cages, with a maximum of five mice per cage. Female B6C3-Tg(APPswe, PSEN1dE9)85Dbo/J) transgenic mice at various ages (3, 5, and 9 months) and their age-matched, non-transgenic littermate controls (wild type, WT) were used in this study. For quality control purposes tail clips were taken at the time of death and a second genotyping was performed to confirm the first.

### ELISA

Hemi-brain samples were analyzed for human Aβ 1-40 and 1-42 using commercially available ELISA kits (Life Technologies: KHB3481 & KHB3441) according to manufacturer’s instruction. Amounts of Aβ were normalized to total protein as determined by BCA assay (Pierce #23225).

### Synaptosome Isolation

All mice were sacrificed at the age indicated. Synaptosomes were isolated as previously described with minor modifications (16). The whole brain was dissected from one mouse and homogenized with 20 strokes in 2 mL of complete sucrose buffer (0.32 M Sucrose; 2 mM EGTA; 2 mM EDTA; 10 mM HEPES (pH 7.4), 1x Protease Inhibitor Cocktail: cOmplete, EDTA-free (Roche 05 056 489 001), 1mM Na_3_VO_4_ (Sodium Vanadate), 1mM Na_4_O_7_P_2_ (Sodium Pyrophosphate). Centrifuged at 800 × g for 10 minutes at 4°C to pellet the membrane fragments and nuclei and collect supernatant (S1) in 15 mL conical tubes then fast frozen in liquid nitrogen and then stored at 70°C until all mouse brain samples for were collected. To minimize technical variability, synaptosomal preparation was performed in age-matched batches. All batch purifications were performed with the same stock buffers within 24 hours of each other. To prepare purified synaptosomes the S1 samples were thawed on ice, centrifuged at 800 x g for 10 minutes at 4°C and collect supernatant (S1’) in high-speed polycarbonate tubes. S1’ fractions were then centrifuged at 10,000 x g for 15 minutes at 4°C to obtain a pellet (P2) containing synaptosomes contaminated with mitochondria and microsomes. P2 was suspended in 500 μl of Complete Sucrose Buffer, vortexed thoroughly then loaded on a sucrose gradient (from bottom to top): 1.18 M – 1.0 M – 0.85 M (all prepared in 10 mM HEPES, pH 7.4, 2 mM EDTA, 2 mM EGTA) prior to centrifuging at 82,500 x g for 1 hour at 4°C. Pure synaptosomes were collected from the interface between 1.0 M and 1.18 M, washed by adding ~3X volume of Complete Sucrose Buffer and centrifuged at 10,000 x g for 15 minutes at 4°C prior to re-suspending in Complete Sucrose Buffer and measuring of protein concentration using a micro BCA Protein Assay (Pierce 22660) according to manufacturer’s instructions. Pure synaptosomes were divided into 50 μg aliquots and stored at -80°C.

### Peptide Preparation

Peptide preparation was performed by the Filter Assisted Sample Preparation (FASP) method as described (17). Briefly, to extract membrane proteins in the synaptosome samples, a 0.3% concentration of Triton X-100 was used to solubilize 50 μg of purified synaptosomes. The sample was then washed with fresh 8 M and 2 M Urea Buffer in a 30 kDa filter (Millipore UFC 503024); the proteins were reduced (10 mM TCEP) and alkylated (25 mM IA) and treated with trypsin (Promega #V5113; 1:100 or 0.05 μg of trypsin per 50 μg sample) overnight in the spin filter. The resulting peptides were desalted in C-18 spin columns (Pierce 89870) according to the manufacturer’s instructions. The peptide concentration was estimated using a NanoDrop 2000 and Bovine Serum Albumin as reference (mass extinction coefficient of 6.7 at 280 nm) resulting in 6-7 μg of peptide per sample. The peptide was then immediately lyophilized for 2 hours, and stored at - 80°C.

### RP-RP MS/MS

The peptides were separated by liquid chromatography using a nanoAcquity UPLC system (Waters) coupled to a LTQ Orbitrap mass spectrometer (Thermo Fisher Scientific). Peptide mixtures (2 μg) were loaded onto a 300-μm × 50-mm XBridge C18, 130-Å, 5-μm column maintained at pH 10.0, eluting peptides in six fractions corresponding to 10%, 15%, 20%, 30%, 40%, and 60% buffer B1 (buffer A1: 20 mM ammonium formate, pH 10.0; and buffer B1: 100% acetonitrile). Steps were eluted from the high pH column at 20 μl/min onto a 180-μm × 20-mm C18, 100-Å, 5-μm trap column, which was then switched in-line with the analytical column and eluted as in the 1D method. For 1D analysis, a BEH C18 reversed phase column (25 cm × 75 μm i.d., 1.7 μm, 100 Å; Waters) was used for the analytical separation using a linear gradient from 90% buffer A2 (0.1% formic acid) to 40% buffer B2 (0.1% formic acid and 80% acetonitrile) over 60 min at a flow rate of 300 nL/min.

MS/MS data were collected by an enabling monoisotopic precursor and charge selection settings. Ions with unassigned charge state were excluded. For each mass spectrometry scan, the 10 most intense ions were targeted with dynamic exclusion 30 s, 1 D exclusion width, and repeat count equal to 1. The maximum injection time for Orbitrap parent scans was 500 ms, allowing 1 microscan and automatic gain control of 106. The maximal injection time for the LTQ MS/MS was 250 ms, with 1 microscan and automatic gain control of 104. The normalized collision energy was 35%, with activation Q of 0.25 for 30 ms.

### Data Analysis

The raw MS/MS data from all samples were analyzed by MaxQuant (37)(version 1.5.2.8). Andromeda (38), a probabilistic search engine incorporated into the MaxQuant framework was used to search the peak list against the Uniprot_MOUSE database (UniProtKB release 2016_098, entries: 82,200). Common contaminants were added to this database. The search included cysteine carbamidomethylation as a fixed modification and N-terminal acetylation and methionine oxidation as variable modifications. The false discovery rate (FDR) was set to 0.01 for both peptide and protein identifications. Enzyme specificity was set to trypsin allowing N-terminal cleavage to proline. Two miscleavages were allowed, and a minimum of seven amino acids per identified peptide were required. Peptide identification was based on a search with an initial mass deviation of the precursor ion (ITMS) of up to 0.5 Da, and the allowed fragment mass deviation (FTMS) was set to 20 ppm. Razor peptides were used for quantification; unmodified or with the modifications specified above. To match identifications across different replicates and adjacent fractions, the “match between runs” option in MaxQuant was enabled within a matching time window of 0.7 min.

Bioinformatics analysis was done with Perseus (39)(version 1.4.1.3) tools available in the MaxQuant environment. The proteins only identified by site, from the reverse database, and contaminant proteins were removed from the matrix. Categorical annotation by age and type was performed resulting in 6 groups: AD_3M, AD_5M, AD_9M, WT_3M, WT_5M, & WT_9M, with an n=3 in each group. The protein matrix was reduced to those identifications with at least one valid observation in each group. A multiple-sample test was run between WT_3M, WT_5M, & WT_9M using a Bengamini-Hochburg FDR=0.25 (7 significant) and these proteins were removed from the matrix. The 9 WT samples were then pooled into one WT group and the data was exported for further analysis with R version 3.3.1 (2016-06-21) and the package limma (21,23) (version 3.28.14). Limma was then used to calculate the log_2_ fold change and the empirical Bayes p-Value between the following groups: AD_9m & WT, AD_5m & WT, AD_3m & WT, WT_9m & WT, WT_5m & WT, and WT_3m & WT.

